# RobustClone: A robust PCA method of tumor clone and evolution inference from single-cell sequencing data

**DOI:** 10.1101/666271

**Authors:** Ziwei Chen, Fuzhou Gong, Liang Ma, Lin Wan

## Abstract

Single-cell sequencing (SCS) data provide unprecedented insights into intratumoral heterogeneity. With SCS, we can better characterize clonal genotypes and build phylogenetic relationships of tumor cells/clones. However, high technical errors bring much noise into the genetic data, thus limiting the application of evolutionary tools in the large reservoir. To recover the low-dimensional subspace of tumor subpopulations from error-prone SCS data in the presence of corrupted and/or missing elements, we developed an efficient computational framework, termed RobustClone, to recover the true genotypes of subclones based on the low-rank matrix factorization method of extended robust principal component analysis (RPCA) and reconstruct the subclonal evolutionary tree. RobustClone is a model-free method, fast and scalable to large-scale datasets. We conducted a set of systematic evaluations on simulated datasets and demonstrated that RobustClone outperforms state-of-the-art methods, both in accuracy and efficiency. We further validated RobustClone on 2 single-cell SNV and 2 single-cell CNV datasets and demonstrated that RobustClone could recover genotype matrix and infer the subclonal evolution tree accurately under various scenarios. In particular, RobustClone revealed the spatial progression patterns of subclonal evolution on the large-scale 10X Genomics scCNV breast cancer dataset. RobustClone software is available at https://github.com/ucasdp/RobustClone.

## 1 Introduction

Tumor evolution has been a subject of longstanding discussion (1). The evolutionary mechanisms underlying cancer progression are believed to guide the principles in understanding, predicting, and controlling cancer progression, metastasis, and therapeutic responses (2). A tumor is comprised of subpopulations of cells with distinct genotypes called subclones (3). In order to understand the genetic intratumoral heterogeneity, computational methods have been developed, taking advantage of high-throughput, next-generation sequencing data (4), to deconvolve subclonal genotypes and/or infer their evolutionary relationships through the reconstruction of subclonal phylogenetic trees (5; 6; 7; 8; 9; 10; 11; 12) from bulk DNA sequencing data of tumor cells.

The rapid advances in single-cell sequencing (SCS) have greatly enhanced the resolution of tumor cell profiling, promising better characterization of intratumoral heterogeneity (3; 13). While pioneering work utilizes single-cell copy number variation (scCNV) profiles to construct tumor cell phylogenies (14; 15), many others work on single-cell single nucleotide variation (scSNV) data with application of traditional phylogenetic methods. (16) and (17) applied distance-based methods, such as UPGMA or neighbor-joining (NJ) (18). Methods based on more complex models, like maximum likelihood or Bayesian (19), have also been applied to infer tumor phylogeny with scSNV data (20; 21).

However, because of technique issues, such as library preparation and dropout events, current SCS data are known to be error-prone, thereby limiting the direct application of traditional phylogenic approaches. Three common types of errors are often associated with SCS data: false positive (FP) and false negative (FN) mutations, as well as missing bases (MBs). FPs and FNs are usually caused by allelic dropout events, a very common problem in SCS in which the mutant allele fails to amplify. The false positive rate (FPR) varies from 0.1 to 0.43, as reported in many studies (15; 16; 22; 23). False positives occur on the order ~ 10^−5^, exceeding the somatic mutation rate (15; 16; 22). MBs may have issues involving sequencing depth, and they occur in low-coverage regions during sequencing. The reported missing rate (MR) can be as high as 58% in single-cell sequencing (22; 23). When these errors in the data occur together, downstream analysis can be biased.

Methods that explicitly account for errors in SCS data, especially scSNV data, have emerged in recent years. SiFit (24) is a likelihood-based approach, which employs a finite site model of evolution and infers cell phylogeny, considering SCS errors. BEAM (25) is a Bayesian method that improves the quality of single-cell sequences by using the intrinsic evolutionary information in the single-cell data in a molecular phylogenetic framework. SCG (26) also models various technical errors by clustering single cells into subclones with a hierarchical Bayesian model and then infering the constituting genotypes. Other methods, such as SCITE (27) and OncoNEM (28), also model the noise of SCS, as well as construction of mutation or subclone trees based on scSNV. Among the above-mentioned methods, each has its own merits, as they all perform acceptably well under the present amount of data size. However, single-cell techniques are rapidly evolving (29), correspondingly lowering the cost of sequencing. In addition, the size of single-cell samples and the number of mutations, which can be used in the analysis, are expected to increase in the very near future (30). These changes could result in a dramatic increase of computational intensity for approaches with likelihood, as well as Bayesian-based, algorithms.

Now, the low-rank matrix factorization method, robust principal component analysis (RPCA), for recovering low-dimensional subspace from corrupted data, is being extensively studied (31; 32; 33; 34). RPCA is a generalization of the standard principal component analysis (PCA) by introducing some robustness. Instead of approximating observation with a low-rank matrix, as in PCA, RPCA approximates observed matrix by decomposing the matrix as the sum of a low-rank matrix plus a sparse component to model the corrupted variables. RPCA works well, even with respect to grossly corrupted observations. The decomposition of RPCA can be implemented by scalable and fast algorithms, such as the Augmented Lagrange Multiplier Method (ALM) (33). RPCA is also naturally extended to model corrupted data in the presence of missing entries (34). RPCA has wide applications in fields such as image processing (34) and bioinformatics (e.g., the imputation of single-cell RNA sequencing data (35)).

In this study, we present a computational framework, RobustClone, allowing the recovery of subclone genotypes based on the GTM of either scSNV or scCNV data and reconstructing the subclonal evolutionary tree. By applying extended RPCA, which tolerates more missing entries, RobustClone first imputes and recovers the cell genotype profiles from error-prone scSNV/scCNV data. It then identifies subclones by clustering cells using the Louvain-Jaccard method (36; 37). Finally, RobustClone reconstructs the subclonal evolutionary tree by finding its minimum spanning tree (MST). By applying RobustClone to simulated and real data, we will demonstrate the power of RobustClone in allowing the identification of real GTM and reconstruction of subclonal evolutionary trees under various scenarios.

## 2 Methods

In this section, we first introduce RPCA and the extended RPCA algorithms, which are used to recover the low-rank subspace from data matrix with corrupted and/or missing entries (Section 2.1); we then describe how the proposed computational framework, termed RobustClone, recovers the true genotype matrix of tumor cells, identifies tumor subclones, and reconstructs subclonal evolutionary trees, all based on tumor single-cell sequencing data (Section 2.2). In Section 2.4 and 2.3, we will introduce the data and method used for application and evaluation of RobustClone.

### 2.1 RPCA and extended RPCA

#### 2.1.1 RPCA

As a popular tool to recover low-rank matrix, standard PCA is based on the assumption that all sample points are drawn from the same statistical or geometric model (34). In practice, however, the entries of data matrix can be corrupted by gross errors, making standard PCA less robust to intrasample outliers (34). In this regard, (31) proposed the robust PCA method (RPCA) to recover low-rank matrix from data with corrupted entries. The RPCA problem can be solved with the Augmented Lagrange Multiplier Method (ALM), a fast and scalable algorithm proposed by (33).

Assume that the data matrix *D*_*m*×*n*_ is generated as the sum of two matrices *D* = *A*_0_ + *E*_0_, where *A*_0_ represents a low-rank data matrix, while *E*_0_ represents the intrasample outliers. We also assume that many entries of *D* are not corrupted, thereby causing many entries of *E*_0_ to be zero. As a consequence, the RPCA problem becomes one of decomposing matrix *D* as the sum of a low-rank matrix *A* and a sparse matrix *E*, satisfying

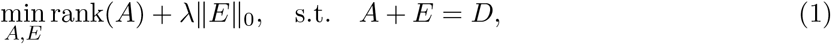

where ∥*E*∥_0_ is the number of nonzero entries in *E*, and λ is the regularization parameter that balances the two terms. However, the task of recovering the low-rank matrix *A* and the sparse signal *E* in Problem (1) is generally NP-hard (34).

To lessen burdens of computation, Problem (1) is convex relaxed, as

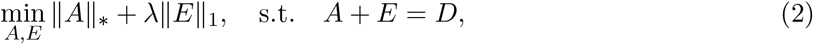

where ∥ · ∥_*_ and ∥ · ∥_1_ denote the nuclear norm and the *ℓ*_1_ norm of matrix, respectively. We call Problem (2) the relaxed version of RPCA. With a probability of almost 1, it has been theoretically validated that the relaxed RPCA can decompose *D* and exactly recover the unknown matrices *A* and *E* under rather broad conditions (see (31; 34)). The optimal choice of λ has been shown to be 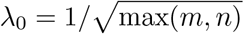 (31; 34).

The constrained optimization problem can be solved by the ALM algorithm proposed by (33), a special case of the alternative direction method of multipliers (ADMM), by applying it on the following augmented Lagrangian function:

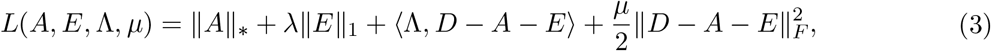

where ∥ · ∥_*F*_ is the Frobenious norm of matrix, Λ is the Lagrange multiplier, 〈·, ·〉 is the inner product, λ and *μ* are the regulation terms. See details of the algorithm in Supplementary Note 1 and Algorithm S1.

#### 2.1.2 Extended RPCA

We further extend RPCA to the cases where entries in the data matrix can be corrupted and/or missing/incomplete. In order to handle the missing entries, we define a linear operator 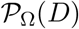, which maps the missing/incomplete entries to 0, while keeping the observed entries, as

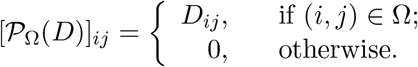

The extended RPCA problem is then formulated as follows (34; 38; 39):

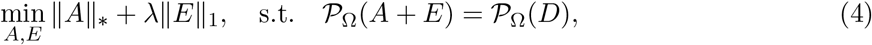

with the intention of recovering the low-rank matrix and the sparse component (*A*, *E*) of *D* = *A*+*E* only from the observations 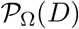. The low-rank and sparse components can be exactly recovered, as well, with high probabilities under conditions similar to those for RPCA (34; 38; 39). (39) show that the optimization problem (4) is equivalent to solving the following constrained optimization problem:

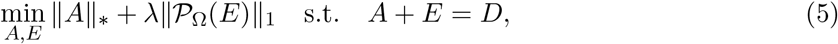

which can be solved by applying the ALM algorithm (Algorithm S2) with the following augmented Lagrangian function:

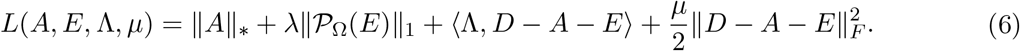

In choices of λ, we put more weight on 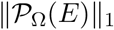 when the data matrix has higher MR, i.e., 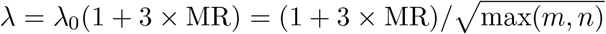. When there are no missing entries (MR = 0), then λ = λ_0_, which is the optimal choice of RPCA.

### 2.2 RobustClone

We next introduce the proposed computational framework, RobustClone, which lays on the foundation of RPCA and extended RPCA. It applies tumor single-cell sequencing data to recover the true genotypes of cells, identify subclones, and reconstruct subclonal evolution trees.

RobustClone takes the input of observed GTM of *Y*_*m*×*n*_ = [*y_ij_*]_*m*×*n*_ from either scSNV or scCNV data, where *y_ij_* denotes the genotype of locus *j* ∈ {1, ⋯, *n*} of individual cell *i* ∈ {1, ⋯, *m*}. The value of *y_ij_* can be either binary (e.g., *i* ∈ {0, 1}: “0”-unmutated, “1”-mutated), or ternary (e.g., *i* ∈ {0, 1, 2}, the number of mutant alleles) for scSNV data, and it can be a nonnegative integer for scCNV data (e.g., *i* ∈ {0, 1, 2, ⋯, *p*}, the number of copies of a DNA fragment, where usually the normal case in the diploid genome has *i* = 2). We use “NA” for entries in the GTM with incomplete or missing values.

The flowchart of RobustClone is shown in Figure 1. The main steps for RobustClone are organized in the following subsections.

**Figure 1:**
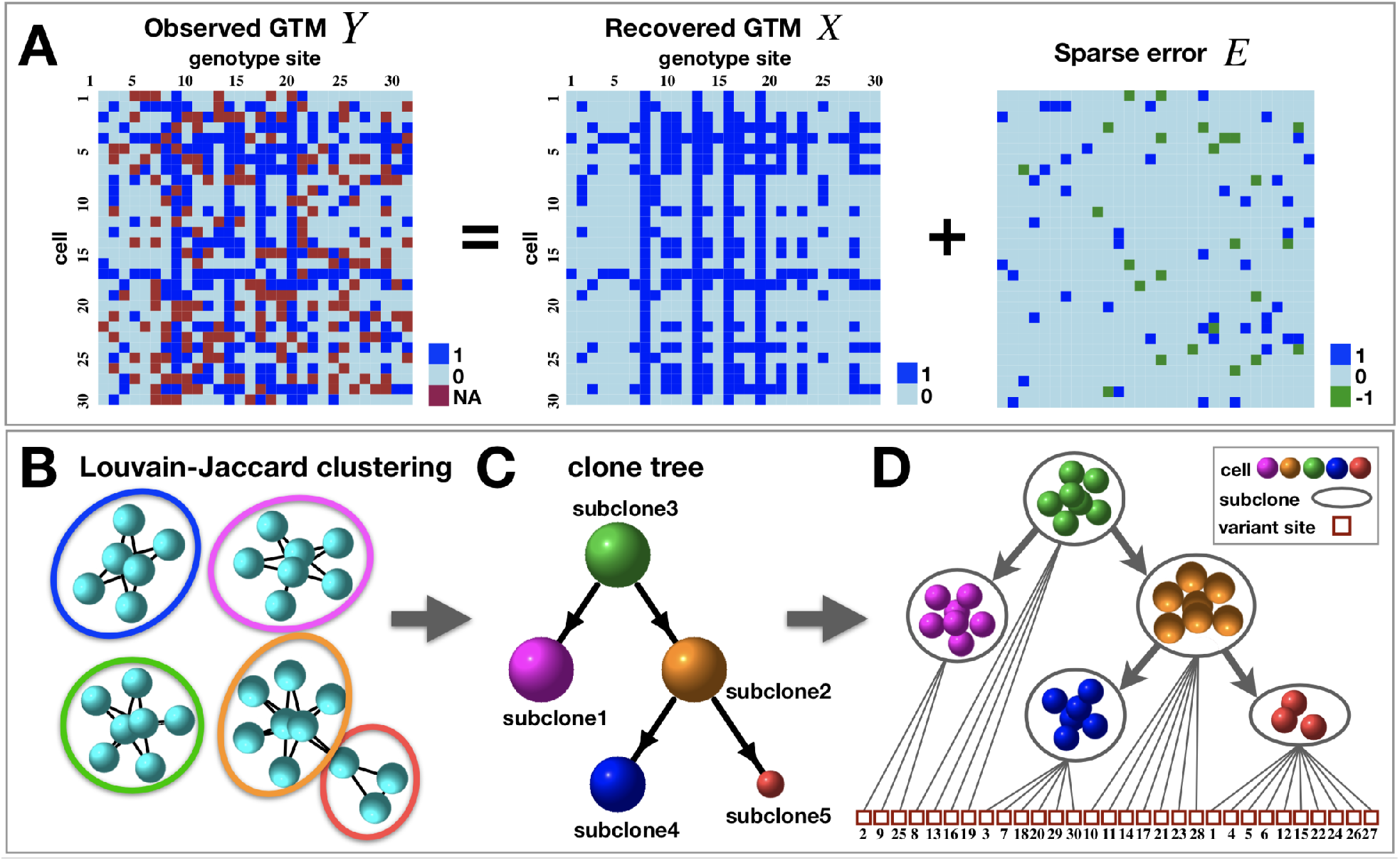
Overview of the computational framework of RobustClone that recovers the true genotypes of cells, identifies subclones, and reconstructs subclonal evolution trees using tumor single-cell sequencing data. **(A)** RobustClone decomposes the observed genotype *Y* into the sum of the lowrank genotype matrix *X* and a sparse matrix *E* by RPCA or the extended RPCA model. **(B)** RobustClone divides the individual cells into clusters, as our identified subclones, by applying the Louvain-Jaccard method on the recovered low-rank genotype matrix *X*. **(C)** RobustClone reconstructs the subclonal evolutionary tree: RobustClone identifies the subclonal tree by finding the minimum spanning tree using Euclidean distance after extracting the consensus subclonal genotypes; the radius sizes of the nodes on the subclonal tree are proportional to the number of cells contained in each subclone. **(D)** The subclonal evolutionary tree that describes the subclonal development of the tumor and the newly mutated genotypes of each subclone from its parent subclone.

#### 2.2.1 Recover the true genotype matrix of cells

Tumor cells are heterogeneous. Nevertheless, as a subpopulation of cells with nearly, or completely, identical genetic composition in subclones, there should be generally fewer subclone types than the number of cells. Besides missing entries, the observed single-cell GTM *Y*_*m*×*n*_ is often incorporated with errors such as FPs and FNs, thus making it extremely difficult to directly identify subclones from *Y*. Therefore, making imputations to the missing entries and correcting the erroneous entries are essential steps.

RobustClone will first recover a matrix *X*, which approximates the underlying true genotypes of cells, from the original observed matrix *Y*. Since tumor cells represent a subpopulation of nearly, or completely, identical cells grouped into subclones, the true genotype matrix would be in low-rank. It is well known that the standard low-rank factorization method, PCA, is sensitive to noise and that it lacks robustness (31). Therefore, we apply the RPCA method to recover the low-dimensional subspace *X* from *Y*. When there are no missing entries in *Y*, we can decompose observed *Y* with corrupted entries, as follows:

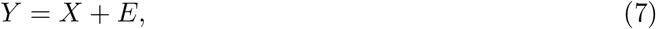

where *X* is a low-rank matrix which approximates the true genotypes of cells, and *E* is a sparse matrix representing noise in the original data. For scSNV data with binary values, we can further denote *E* = *E*_1_ + *E*_2_, where *E*_1_ represents the noise caused by FPs, and *E*_2_ represents the noise caused by FNs. For scCNV data, *E* can be the noise generated in DNA sequencing and/or the errors caused during the estimation of copy numbers.

Under this scheme, RobustClone directly applies the RPCA algorithm (Problem 2 in Section 2.1.1) to recover the low-rank matrix *X*.

Generally, dropout events frequently happen under current single-cell sequencing technologies. This means that only a fraction of the DNA present in each cell is being sequenced. The dropout frequency can reach as high as 58% in a single dataset of scSNV (22; 23). These frequent dropout events can lead to serious incompletion in the observed GTM of *Y*. To handle this issue, we borrow low-rank matrix completion techniques and extended RPCA (Section 2.1.2) to recover the low-rank matrix *X* and sparse matrix *E* under the constraints of *Y* = *X* + *E*, but only on the observed set, i.e., 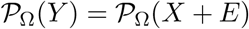, where Ω is the observed set (see the definition of linear operation 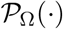 in Section 2.1.2).

Therefore, in the presence of missing entries in *Y*, RobustClone applies the extended RPCA model (5) to recover low-rank and sparse components for incomplete and grossly corrupted SCS data simultaneously (Figure 1A).

#### 2.2.2 Identify subclones by Louvain-Jaccard clustering method

RobustClone divides individual cells into clusters, in a manner similar to that of our identified subclones, by applying the Louvain-Jaccard method (36; 37) on the imputed and recovered lowrank genotype matrix *X*. Matrix *X*, which is solved by the convex relaxation method based on nuclear norm of matrix instead of its rank, is clear and a good approximation of the true genotype. However, it still cannot guarantee an error-free state. Thus, in real applications, we cannot identify subclones simply by aggregating the identical rows of *X*.

To cluster single cells, RobustClone adopts the Louvain-Jaccard method (36; 37), which has wide applications in the clustering of single-cell RNA sequencing data (40) (Figure 1B). The Louvain-Jaccard method is a network-based fast community detection algorithm. The community detection algorithm is based on the idea of modularity, as described by (41), the rationale being that nodes in the same community have more edges than nodes between communities. In its implementation, the Louvain-Jaccard method first constructs a *k*-nearest neighbor (k-NN) graph of *m* cells based on Euclidean distance. The choice of parameter *k* is empirically dependent on the sample size (number of single cells), and we demonstrate that the results by Louvain-Jaccard algorithm is robust to the choices of *k* (see Supplementary Note 2 and Figures S8, S9, S10).

However, we want to emphasize that RobustClone is not restricted to the Louvain-Jaccard algorithm. Clustering methods, such as hierarchical clustering and K-means algorithm, can also be adopted by RobustClone to identify subclones. The Louvain-Jaccard algorithm does not require pre-specification of the number of subclones, and it is a fast and scalable algorithm that can be applied to large-scale datasets.

#### 2.2.3 Reconstruct subclonal evolution tree

RobustClone reconstructs the subclonal evolution tree of tumor by the following two steps.

First, the consensus genotypes are extracted for each subclone identified in Section 2.2.2. Since the cells within the same subclone identified by the Louvain-Jaccard method are homogeneous in genotypes, with their genotypes being identical, or almost identical, with minor errors, RobustClone simply extracts the consensus genotypes for each genetic site: 1) for scSNV data, the genotype with highest frequency among the cells from the same subclone is selected as the consensus, and 2) for scCNV data, the median of the copy numbers among the cells from the same subclone is taken.

Second, the subclonal evolutionary tree is reconstructed (Figure 1CD). RobustClone calculates the Euclidean distance between each pair of subclones using the consensus genotype vector and then finds the minimum spanning tree (MST) among the subclones based on Euclidean distance. To determine the root of the tree, RobustClone selects the node with the shortest Euclidean distance to a given reference/normal sample.

### 2.3 Evaluations

We compared the performance of RobustClone to BEAN, SCG and SiFit (24; 25; 26) under various simulated scenarios. The performance of each was evaluated on basis of several metrics that measure different aspects of the goodness of the recovered GTM, including 1) the *FPs*+*FNs* ratios of output GTM to input GTM, 2) the percentage of correctly imputed missing bases and 3) the error rate of the recovered GTM to the ground truth. Metrics 1) and 2) were also utilized by (25).

In addition, we also evaluated the performance of each on the reconstructed phylogenetic tree by using Robinson-Foulds (RF) distance (42; 43), which measures the disparity between two phylogenetic trees. RF distance is calculated between the reconstructed tree from the recovered GTM (e.g., RobustClone and SCG) or the output tree (e.g., BEAM and SiFit) and the tree constructed on the ground truth GTM (e.g., Maximum Likelihood tree).

### 2.4 Data

#### 2.4.1 Simulation Data

In the simulation study, we adopted the two-step simulation procedure used in (28) (see details in Supplementary Note 3), which first generates a clonal lineage tree with a given number of clones and then simulates the corresponding genotypes. To systematically evaluate RobustClone and compare it with other methods, we simulated 25 datasets with various parameters (hereinafter denoted as comparison datasets). We fixed the number of cells for all comparison datasets to 1000. We divided the 25 simulated comparison datasets into 5 groups (denoted as G1-G5). Each group contained 5 datasets which were simulated by changing the setting in one of the 5 parameters (e.g., the number of genetic sites (loci) (#GS), FPR, FNR, MR, and the number of simulated subclones (#SC)), while keeping the other 4 fixed (see Table S1 for details).

To evaluate the efficiency and accuracy of RobustClone on different data size, we simulated 3 datasets of GTMs, respectively, with the sizes of cells × genotypes as 5000 × 1000 (constituting 100 subclones), 500 × 500 (constituting 5 subclones) and 50 × 100 (constituting 3 subclones). The other parameters were identical for the 3 simulations, which were set as follows: FPs (15%), FNs (15%) and MBs (20%).

#### 2.4.2 Real Data

We also tested RobustClone on 4 real datasets, including 2 each of scSNV and scCNV datasets. The scSNV datasets were 1) high-grade serous ovarian cancer (HGSOC) data (26; 44) and 2) essential thrombocythemia (ET) data (22). The 2 scCNV datasets were 1) xenograft breast tumor data (hereinafter denoted as SA501X3F) (45; 46) and 2) breast cancer data from 10X Genomics (https://www.10xgenomics.com/solutions/single-cell-cnv/).

## 3 Results

### 3.1 RobustClone recovers genotype matrix with high accuracy and efficiency on simulation datasets

In order to demonstrate the steps of RobustClone, we generated an illustrative dataset with 5 subclones, 1000 cells and 300 SNV sites, with the settings of other parameters as MR 20%, FPR 15% and FNR 15%, respectively. The result showed that RobustClone could recover the true GTM with high accuracy and that the inferred subclonal tree was consistent with the simulated topology (see details in Supplementary Note 4 and Figure S1).

In a more systematic evaluation, we compared RobustClone with other methods (BEAM, SCG and SiFit) on the 25 simulated comparison datasets (Figure 2). The implementation of SCG for this analysis was based on the genotype-aware D_SCG3 model. Since OncoNEM cannot handle data with a large number of cells, we excluded it in the comparisons.

**Figure 2:**
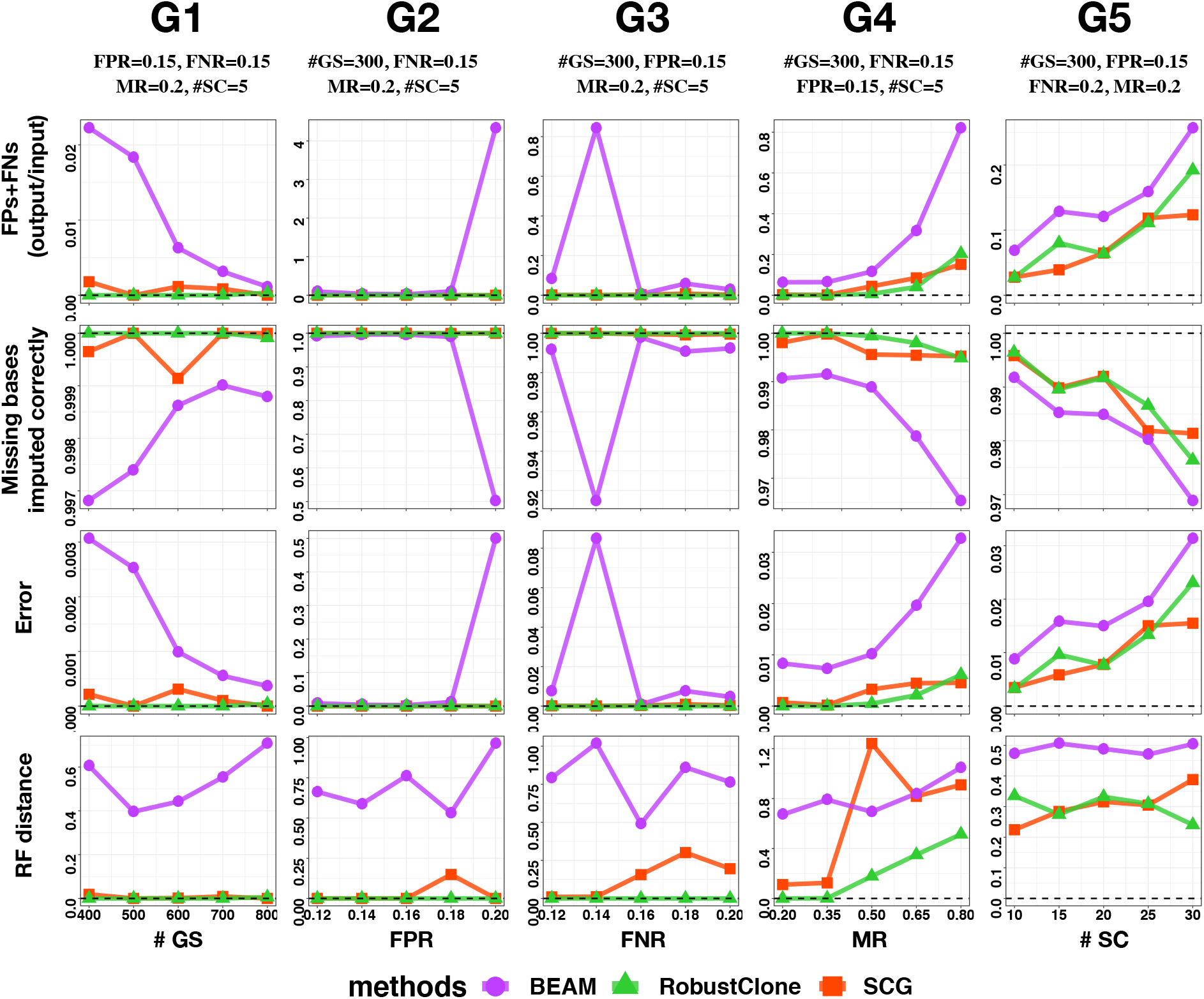
Comparison of accuracy among RobustClone, BEAM and SCG algorithms on the comparison datasets. The comparison datasets were divided into 5 groups (denoted as G1-G5). Each of the 5 groups contained 5 datasets simulated by varying one of the 5 simulation parameters (e.g., the number of genotype sites (#GS), FPR, FNR, MR, and the number of simulated subclones (#SC)), while keeping the remaining 4 fixed. The columns represent G1-G5; the rows are metrics of accuracy by 1) the *FPs* + *FNs* ratios of output GTM to input GTM, 2) the percentage of correctly imputed missing bases, 3) the error rate of the recovered GTM compared to the ground truth, and 4) Robinson-Foulds (RF) distances. In all panels, the position of the black dashed line represents the value of the optimal indicator.

In general, all methods, except SiFit, performed comparably (Figure S2). When increasing the number of actual subclones (SC) or missing rate (MR), all methods showed a specific increase in error rate associated with more output FPs+FNs and less correctly imputed missing bases (Figure 2 G4, G5). BEAM had a steeper increase in error rate when missing bases reached 50% (Figure 2 G4). The increase of false positive rates (FPR) or false negative rates (FNR) in simulated data did not have a significant effect on output error rate of RobustClone and SCG, whereas BEAM was more sensitive to large input FPR (Figure 2 G2, G3). RobustClone not only showed stable and excellent performance as the number of genetic sites (GS) increased in the GTM, but also had the shortest computational time among the compared methods (Figure 2 G1, Table S2). The performance of SCG and BEAM tended to be better with more GS, while SCG had more computational efficiency comparable to that of BEAM. In contrast, SiFit had more errors with larger size of GS (Figure S2).

We further evaluated the computational time and performance of RobustClone on 3 simulated datasets of different size (Table S3). RobustClone performed robustly in small- and medium-sized data with fast computation and low error rate. For data size as large as 5000 × 1000 (cells × GS), RobustClone only took 4.5 minutes of computational time, resulting in an output error rate of 0.015. Although SCG had a close output error rate, its computational time was 27 times greater than that of RobustClone, which took 122 minutes. While BEAM needed 169 minutes in computation time, SiFit failed in data with such size.

In sum, RobustClone showed excellent performance in all simulated scenarios with efficient computational time and low output error rate. The performance and efficiency of SCG was comparable to that of RobustClone and slightly better than BEAM. It significantly outperformed SiFit. It is worth noting that SCG, as well as the other compared methods, was restricted to scSNV data, while RobustClone could also be applied to scCNV data.

### 3.2 RobustClone works robustly on real data with high missing rate

We applied RobustClone to a set of real scSNV data with high missing rate (22). Single-cell exome sequencing from a sample of JAK2-negative myeloproliferative neoplasm (essential thrombocythemia, ET) contains 58 single cells (22) with 712 somatic single nucleotide variants. We applied the binarized GTM preprocessed in (28) as the raw input for RobustClone. The missing rate of this dataset reaches 58% (Figure S3A). The RobustClone algorithm recovered the GTM by imputation of missing entries and correction of erroneous entries (Figure S3B). The 58 tumor cells were then clustered by RobustClone into 3 subclones, respectively containing 25, 19 and 14 cells (Figure S3B). With subclone3 identified as the root, RobustClone found an MST in linear topology that connected all 3 subclones (Figure S3C). This result is consistent with the previous findings in (22; 27; 28).

In addition to scSNV data, RobustClone can also be used to detect copy number heterogeneity and identify clones with scCNV data. To demonstrate this, we applied RobustClone on copy number profiles of cells from the passages of a patient-derived primary triple-negative breast cancer (TNBC) xenograft (SA501X3F data), which contains 260 cells and 20651 genomic bins of copy number states (Figure S4A). RobustClone recovered a GTM with cells clustered into two subclones, where one subclone consisted of 214 cells (denoted as subcloneA), and the other consisted of 46 cells (denoted as subcloneB) (Figure S4B). The copy number profiles of the two subclones are shown in Figure S4C. It can be seen that the difference in copy numbers between subcloneA and subcloneB is mainly presented on the X chromosome (Figure S4B,C). SubcloneA is completely consistent with the major subclone identified in (45; 46; 47). If we take into account the high noise in the data together with the small size of subcloneB and then tune parameter λ to panelize more on the sparsity of the error entries (*E* in (2.1)), we can identify an extra subclone separate from original subcloneB (Figure S5), which is consistent with the clonal subpopulations identified in (45). However, since (45) did not explicitly correct for noise in the GTM before clonal identification, the two derived subpopulations from subcloneB had much uncertainty (46). We believe that the two subclones that resulted from the default setting of RobustClone are more robust.

In order to further assess the robustness of RobustClone on the SA501X3F dataset, we randomly performed a 30% dropout of entries in the original GTM (Figure S6A) and reanalyse it. RobustClone still identified 2 subclones, and the copy number profile of cells in different subclones was consistent with the result in Figure S4B (Figure S6B). We continued to increase the dropout rate to 50% (Figure S6C). RobustClone found 4 subclones containing cells in this order: 213, 45, 1 and 1 (Figure S6D). The two extra subclones each contain 1 cell only, which is separate from one of the original major subclones.

These results indicate that RobustClone is highly robust for scSNV and scCNV data with high missing entries.

### 3.3 Recovering scSNV genotype and inferring subclonal tree of high-grade serous ovarian cancer

We performed RobustClone on a set of high-grade serous ovarian cancer (HGSOC) data (26; 44). The original data matrix contains 420 cells and 43 selected SNV sites with 10.7% missing entries (Figure 3A). RobustClone efficiently recovered the cellular GTM by imputing the missing values and correcting noisy entries in the observed data (Figure 3B). Based on the corrected GTM, RobustClone identified 5 subclones. We labeled them subclone1-subclone5, according to their sizes, respectively consisting of 122, 81, 81, 69 and 67 cells (Figure 3C). As subclone4 had the minimum number of mutations, it was assigned as the root subclone. An MST connecting all five subclones was then constructed by RobustClone based on pairwise clonal distance (Figure 3C). The newly arisen mutations of each subclone, following the topology of MST, could thus be identified (Figure S7A).

**Figure 3:**
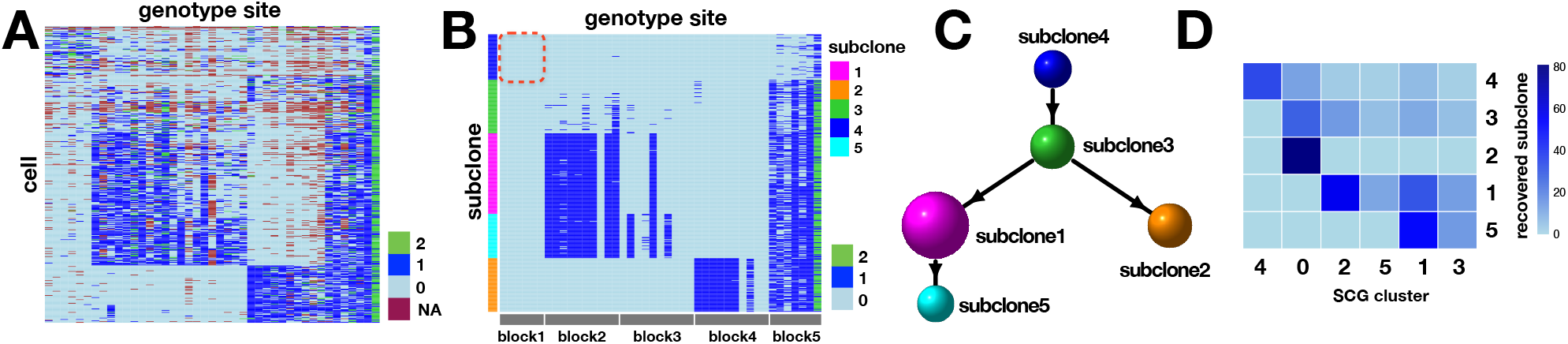
RobustClone reconstructs the subclonal evolution tree on HGSOC data. **(A)** The heatmap of the noisy SNV profile before inference. Each row of the heatmap represents a single cell, and the column represents the GS. **(B)** The heatmap of the GTM recovered by RobustClone. Each row of the heatmap represents a single cell, and the column represents the GS. The genotype sites in the red circle are unmutated, but they were recovered as mutated sites by SCG, as shown in Figure 3b of (26). **(C)** The subclonal evolution tree reconstructed by RobustClone using MST. **(D)** The number of overlapped cells of HGSOC data contained in subclones identified by RobustClone and cells contained in clones identified by SCG.

To better understand the composition and the relationship among subclones, we divided the SNV sites into 5 major blocks. Only subclone4 had some mutations in block5 where site *TP53* mainly presented in heterozygous genotype. The mutations in block5 were carried through all subsequent subclones. These subclones had a high rate of homozygous *TP53* mutant alleles, which is a strong indicator of cancer (48). Subclone3 descended from subclone4 and accumulated sparse mutations in all blocks except block1. The mutations in block2 and 4 define the divergence of subclone1 and subclone2 from subclone3. The smallest of the five blocks, subclone5, which carried more mutations in block3, was derived from subclone1.

We compared the subclones identified by RobustClone to the result of SCG, which identified 6 subclones (clusters) based on the same HGSOC data (26; 44). SCG cluster0 mainly consists of cells in subclone2 and subclone3. The cells in SCG clusters 1, 2, 3 and 5 are mainly distributed in subclone1 and subclone5 (Figure 3D). Subclone4 contained all cells in SCG cluster4 and was thus interpreted to be normal cells. Interestingly, heterozygous and/or even homozygous mutations were recovered by SCG in SNV sites corresponding to block1, but only in cluster4. These precancerous mutations were expected to become “public”, or at least be abundant, in subsequent subclones (49; 50). In this sense, the recovery of GTM by RobustClone with no mutations in block1 in all cells seems to be more reasonable. In addition to the spanning tree of subclones, the subclonal genotypes and/or the corrected GTM could also be used to directly reconstruct the clonal and/or cellular phylogenies by applying any readily available off-the-shelf methods in classic phylogenetics (Figure S7BC) (18).

### 3.4 Revealing the spatial heterogeneity of breast cancer with multi-section large-scale scCNV dataset

We applied RobustClone to large-scale breast cancer scCNV data from 10X Genomics. The frozen breast tissues were from three negative ductal carcinoma *in situ* specimens, which were divided into 5 spatially consecutive parts, denoted as Sections from A to E (Figure 4A). The raw scCNV sequencing data were preprocessed by Cell Ranger pipeline, resulting in a 9050 × 55572 GTM, which respectively included 2061, 2046, 1448, 1665, and 1830 cells from Sections A to E. Each cell was characterized by the ploidy states of 55572 20-kb bins over 7 chromosomes, which covered chr3, chr4, chr5, chr6, chr7, chr8 and chr10, respectively.

**Figure 4:**
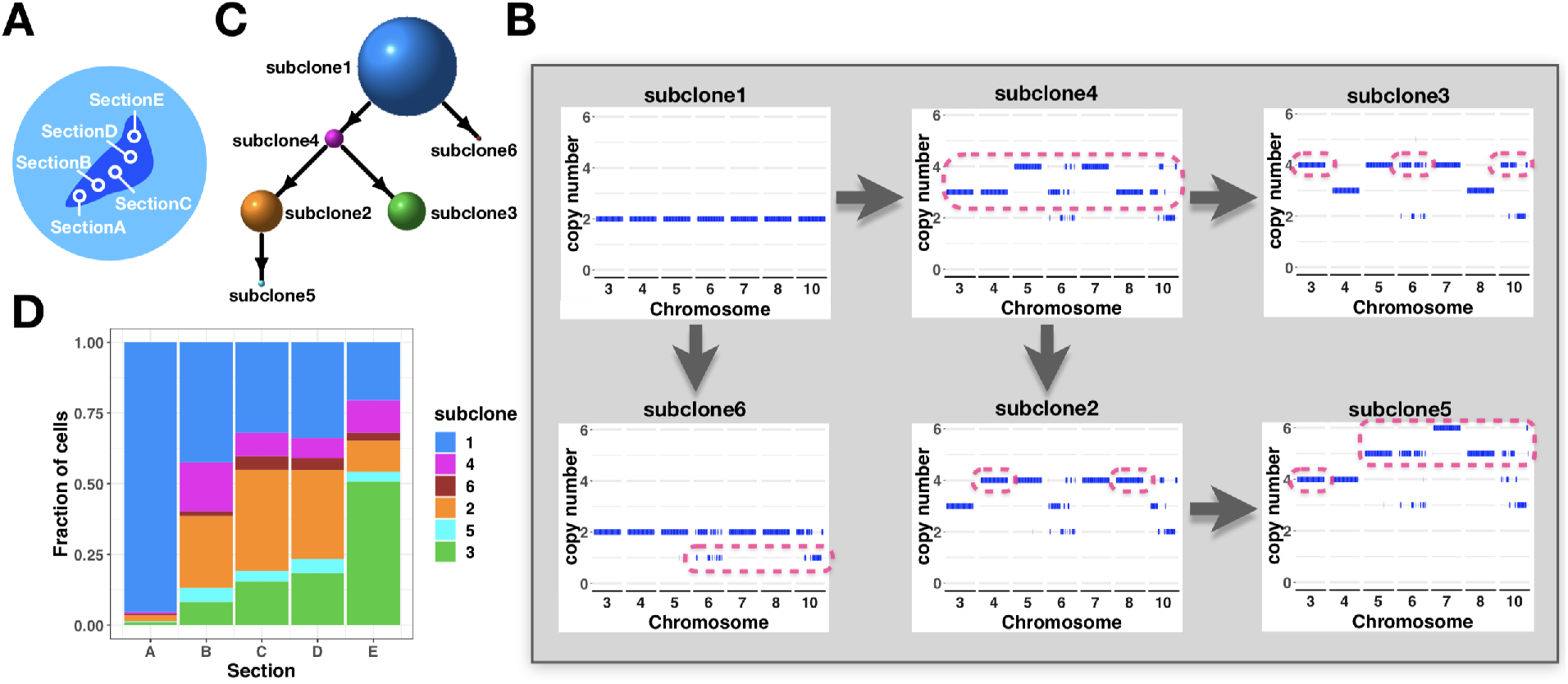
RobustClone reveals a spatial progression pattern of subclones on the large-scale copy number profile of breast cancer dataset from 10X Genomics. **(A)** Schematic representation of the regional division of frozen breast cancer samples; the samples are from the 5 connective spatial points of Sections A-E. **(B)** The copy number profile of subclones recovered by RobustClone. **(C)** The subclonal evolution tree reconstructed by RobustClone. **(D)** Stacked histogram representing subclonal proportions across the 5 sections.

RobustClone first recovered a low-rank scCNV GTM and then identified 6 subclones with 4234, 1808, 1647, 821, 308, and 232 cells, respectively (Figure 4C). The copy number profile (Figure 4B) of each subclone was obtained by taking the median copy numbers of each bin for cells within the same subclone. As the copy number profile of subclone1 was consistently diploid (Figure 4B), it was presumed to consist of normal cells and was, therefore, assigned as the root. RobustClone then found the MST of subclones (Figure 4C). We used red boxes in Figure 4B to highlight the changes between the copy number profiles of each subclone and its parent. Subclone4 diverged from subclone1 and had a gain in copy number in nearly all 7 chromosomes where chromosome 5 and 7 changed from diploid to tetraploid and other chromosomes changed to triploid. Subclone6 also derived from subclone1 with copy number loss in chr6 and 10. Subclone4 further differentiated into two major subclones: subclone2, which changed from triploid to tetraploid on chr4 and 8; subclone3, which gained one more ploidy, mainly on chr3, 6 and 10. Subclone5 gained further copy numbers on all chromosomes, except chr4, on the basis of subclone2.

The subclonal composition of the 5 spatial sections is shown in Figure 4D. Section A is dominated by normal cells of subclone1. Subclone2 occupies the largest proportion of subclones apart from normal cells (subclone1) in the middle sections B, C and D. In contrast, Section E is governed by subclone3. These results reveal the great spatial heterogeneity within tumor.

## 4 Discussion

In this study, we proposed RobustClone, a tool for the robust recovery of noisy scSNV and scCNV data based on the RPCA and extended RPCA. RobustClone is a model-free approach, which achieves high accuracy in the imputation of MBs and correction of FPs and FNs.

The RPCA-based algorithm was applied to the scSMD method for the imputation of singlecell RNA sequencing data (35). Their model assumes that dropout events should be relatively sparse in the original gene expression matrix (35). In the case of the single-cell DNA genotype matrix, however, missing entries may reach as high as 58% (22). Thus, we proposed and applied the extended RPCA in RobustClone, which relaxed the sparsity assumption. By simulation study, RobustClone could achieve high accuracy, even when MR reached 0.8.

Understanding intratumoral heterogeneity and inferring of clonal evolution have long been subjects of research interest (2). Two methods can generally be used to infer clonal relationship with single-cell data. One is jointly modeling errors and clonal phylogeny under a Bayesian or likelihood framework. The other is to first correct the errors in the original single-cell genotype matrix and then construct a clonal tree with the recovered GTM. BEAM, SCITE and SiFit belong to the first kind, and they have good performance when cell numbers or SNV sites are not so large. RobustClone and SCG belong to the latter type. They both exhibit computational efficiency in a large dataset, and they may also utilize the large reservoir of methods in molecular phylogenetics for clonal tree reconstruction. Moreover, RobustClone provides a way to connect subclones to MST, which seems to be more reasonable based on the evolutionary continuity nature of clones. In our comparative studies, RobustClone outperformed the others in both accuracy and efficiency. In addition, unlike other methods that apply only to one type of data, RobustClone can be performed on both scSNV and scCNV data. The application of real data in both cases demonstrated that RobustClone has considerable power in recovering the genotypes of single cells and/or clones, as well as reconstructing cell and/or clone trees.

## 5 Acknowledgements

This work is supported by the National Key R&D Program of China under Grant 2018YFB0704304, NSFC grants (Nos.11571349, 91630314, 81673833), the Strategic Priority Research Program of CAS (XDB13050000), NCMIS of CAS, LSC of CAS, and the Youth Innovation Promotion Association of CAS.

